# Phages indirectly maintain plant pathogen defense through regulation of the commensal microbiome

**DOI:** 10.1101/2024.04.23.590639

**Authors:** Reena Debray, Asa Conover, Britt Koskella

## Abstract

Many infectious diseases are associated with altered communities of bacteriophage viruses (phages). As parasites of bacteria, phages can regulate microbiome diversity and composition and may therefore affect disease susceptibility. Yet observational studies alone do not allow us to determine whether altered phage profiles are a contributor to disease risk, a response to infection, or simply an indicator of dysbiosis. To address this question, we used size-selective filtration to separate plant-associated microbial communities from their respective phages, then transplanted them together or separately onto tomato plants that we subsequently challenged with the bacterial pathogen *Pseudomonas syringae*. Microbial and phage communities together were more disease-protective than either component was alone, an effect that could not be explained by direct effects of phages on either *P. syringae* or the plant host. Moreover, the protective effect of phages was strongest when microbial and phage communities were isolated from neighboring field locations (allopatric phages), rather than from the same host plant (sympatric phages). This suggests a Goldilocks effect in which moderate rates of phage lysis maintain a microbiome community structure that is most resistant to pathogen invasion. Overall, our results support the idea that phage communities contribute to plant defenses by modulating the microbiome.

## INTRODUCTION

Viruses that infect bacteria, or phages, are abundant across habitats and shape microbial ecosystems in significant ways (1–3). For example, when phages replicate within bacterial cells and lyse cells from the inside out, they release the contents of their host cells into the surrounding environment. Through this process, aquatic phages are estimated to release the sequestered nutrients of 10-20% of the ocean’s bacterial microbiome on a daily basis (4). Another key property of phages is their host specificity; a given phage population is typically restricted to a single species of bacteria or even a subset of bacterial strains within a species (5). This means that different bacterial species in a community may experience different levels of phage pressure depending on the composition of the resident phages. In this manner, phages can alter competition outcomes between bacterial species by reducing the growth rates of bacteria that would otherwise dominate (6–8).

Some phages infect and kill bacteria that are pathogenic to plants or animals. These phages are thought to play a role in innate immunity, as mammalian tissues are naturally enriched for phages at potential points of pathogen entry (9,10). Pathogen-targeting phages have long been recognized as an opportunity for managing human, livestock, and crop diseases due to their specificity and their potential to self-replicate and co-evolve with resistant bacteria (11–13).

In comparison, phages that do not directly target bacterial pathogens have received substantially less attention. However, many diseases of animals and plants are characterized by altered phage communities (14–17), suggesting that they may also modulate disease risk. As with the past efforts of the microbiome field to link bacterial community composition to disease phenotypes, it can be difficult to tell from observational data alone whether an altered community is a cause, a consequence, or simply an indicator of a disease state (18,19). Experiments that directly manipulate phage presence and/or composition are therefore needed to identify whether and how phages shape disease susceptibility.

A relationship between phage activity and disease could arise through several pathways that are not mutually exclusive (Figure **1**). One possibility is that direct recognition of phages by the immune system of the plant or animal host alters immune defenses against pathogens. Phages share some molecular features with mammalian viruses (20) and have even been shown to promote antiviral cytokine production in humans and mice (21,22). In one case, a filamentous phage of *Pseudomonas aeruginosa* triggered an antiviral pathway that suppressed phagocytosis of bacterial cells, exacerbating the *P. aeruginosa* infection (23). Another possibility is that phage-mediated lysis of bacteria triggers an immune response to the contents of the bacterial cells. For example, lysis can release lipopolysaccharide, a component of the bacterial membrane that is recognized by plant and animal immune systems (24–27).

**Figure 1.**
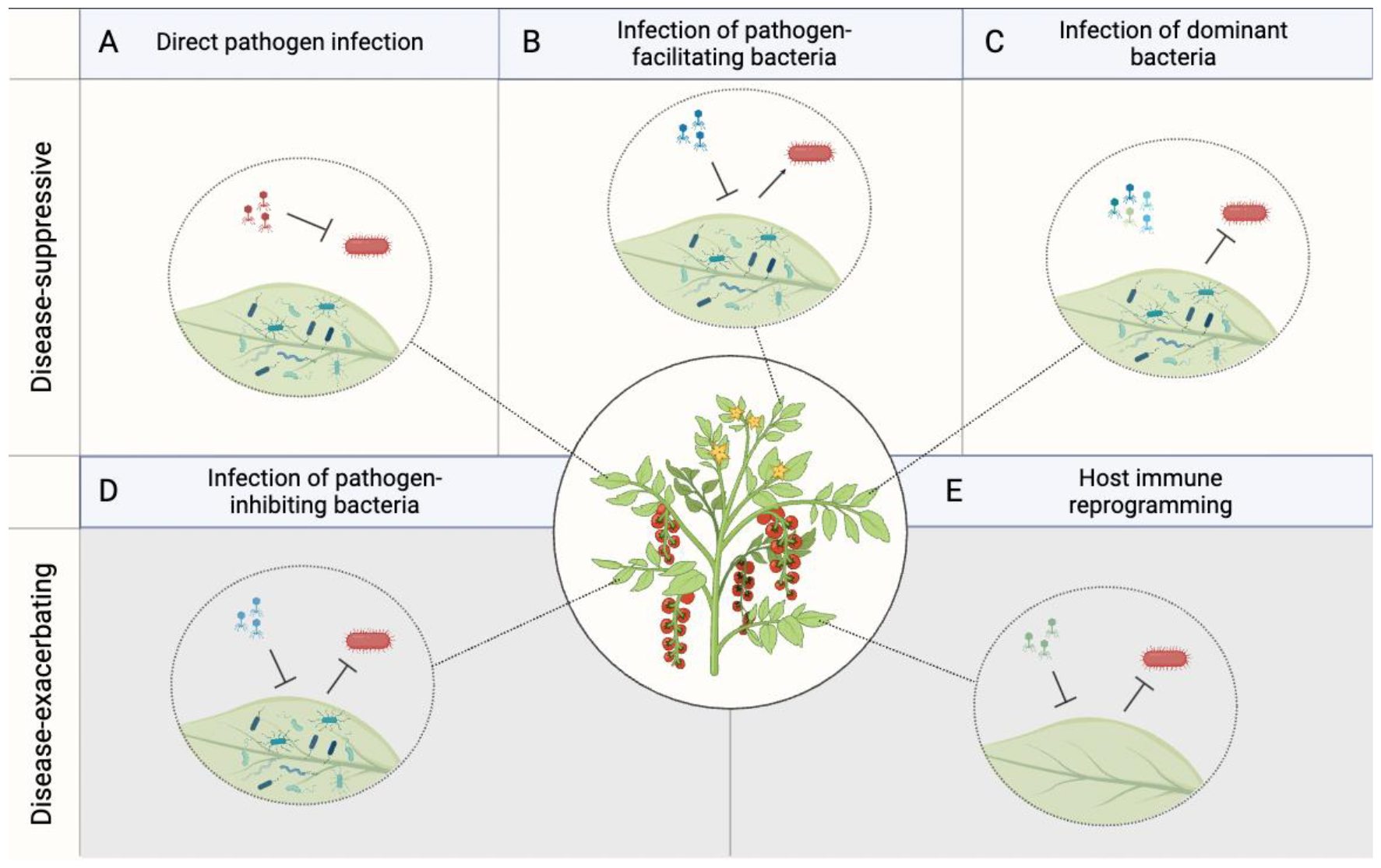
Possible mechanisms of phage modulation of pathogen infection. Phages may suppress disease by **(a)** directly reducing pathogen population sizes (11–13,49–51), **(b)** reducing the population sizes of microbiome members that facilitate pathogen infection, or **(c)** restricting dominant bacteria and in turn facilitating other protective bacterial species that would have otherwise gone extinct. Phages may facilitate disease by **d)** reducing the population sizes of protective microbiome members (16), or **e)** reprogramming host immunity to trigger an antiviral response and suppress the antibacterial response (currently documented only in mammalian hosts, (23)).

Phages may also influence disease outcomes by altering the composition of the resident microbiome. The ability of microbial pathogens to establish in hosts and cause disease symptoms often depends on interactions with other microbiome members. For instance, resident bacteria with similar ecological niches to pathogens can limit pathogen colonization through competition (28,29). Other microbiome members can activate or suppress plant defenses (30–32) or produce compounds that directly inhibit pathogens (33,34). The resident phage community may therefore influence disease susceptibility by altering the abundances of pathogen-facilitating or pathogen-inhibiting bacteria. For example, variation in the severity of bacterial wilt disease in the tomato rhizosphere was attributed to the presence of pathogen-inhibiting bacteria and their associated phages. Tomato plants inoculated with inhibitory bacteria were more resistant to bacterial wilt, but this protective effect vanished in the presence of inhibitor-associated phages (16).

A particularly important feature of phages for disease is their strain- or species-specific killing ability. Ecological models predict that any surge in the abundance of a bacterial species is followed by a surge in its specialized phages, which then reduce the population size of the formerly abundant bacteria (35–39). By restricting dominant bacteria, phages may open niches for other, possibly protective, bacteria that would have otherwise been rare or driven extinct. In support of this hypothesis, bacteria in several habitats form more species-rich and synchronous communities when combined with a diverse set of phages than in the absence of phages (40–42).

In this study, we asked whether phages can modulate plant disease susceptibility, and if so, whether they do so by directly infecting pathogenic bacteria, by stimulating the plant immune system, or by reshaping the commensal microbiome. We collected leaves from field-grown tomato plants and then separated their bacterial and viral microbiomes using size-selective filtration. We transplanted the bacterial and viral communities either together or separately onto tomato leaves in a series of growth chamber experiments. This approach is analogous to predator exclusion experiments that have been conducted in macro-scale ecosystems to determine how top-down processes shape community structure (43,44). We then challenged the plants with *Pseudomonas syringae*, a generalist bacterial pathogen that infects nearly 200 plant species, including a number of economically important crops such as wheat, barley, soybean, kiwifruit, and tomato (45,46). Previous research shows that applying a supplemental microbiome to tomato leaves can limit subsequent *P. syringae* growth and symptom progression (34,47,48), indicating a potential role for the phyllosphere microbiome in disease susceptibility. We find that this protective effect is attenuated in the absence of the phage community. We further show that the contribution of phages to disease protectiveness could not be explained by phage interactions with the plant immune system or by direct effects of phages on *P. syringae* alone. Rather, the effect of phages was contingent on the presence and identity of their bacterial hosts.

## RESULTS

### Phage depletion reduces microbiome-mediated protection

To assess the role that the phage component of the microbiome plays in resistance to bacterial infection, we sampled epiphytic microbial communities from field-grown tomato plants and used a series of membrane filtration and concentration steps to separate cellular microbes from their associated phages. We then transplanted microbial communities, with or without their respective phages, onto juvenile tomato plants in the lab (see **Table 1** for a summary of experiments performed). We subsequently challenged plants with the bacterial pathogen *Pseudomonas syringae*. We measured *P. syringae* population sizes within leaf tissue after 24 hours, as this metric can be accurately quantified with droplet digital PCR (52) and correlates with disease epidemiology and plant susceptibility to frost damage (53). *P. syringae* pv. tomato (strain PT-23) population sizes were approximately 35% lower in plants treated with the ‘microbiome + phages’ inocula than in control plants that were not treated with any supplemental microbiome or phages prior to infection (t = 2.11, df = 17, p = 0.0496, Figure **2a**). Neither the ‘microbiome only’ or the ‘phages only’ components were individually protective against *P. syringae*, suggesting that this effect depended on interactions between phages and their hosts.

**Table 1.**
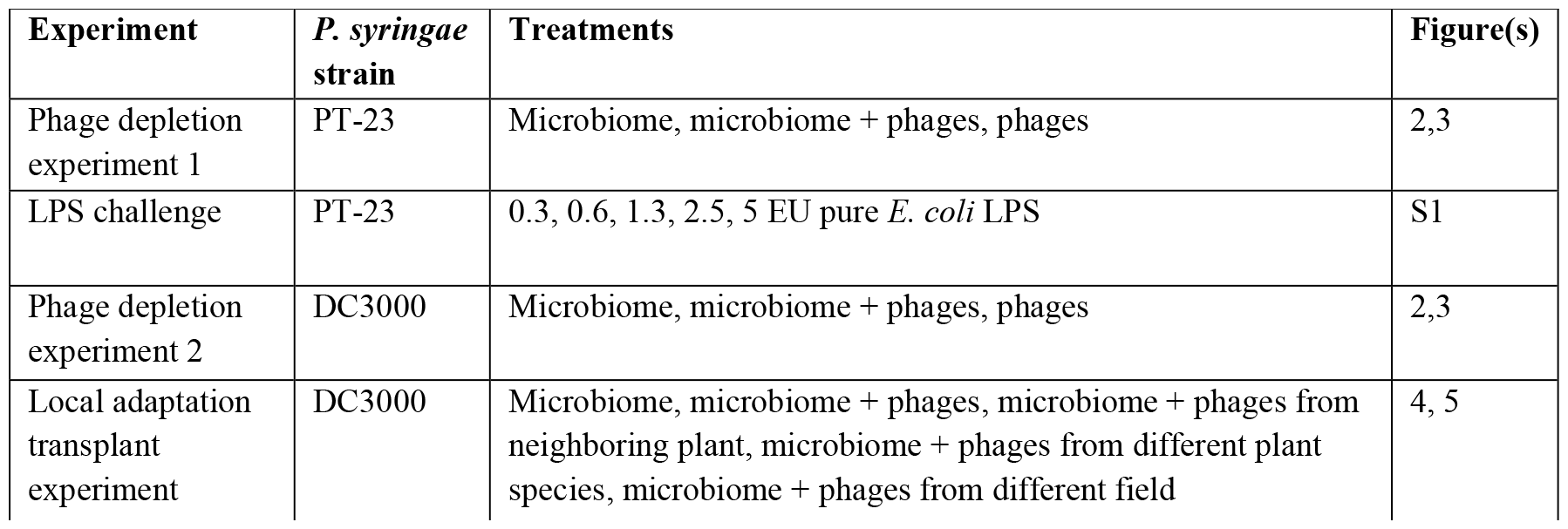
Summary of experiments performed.

| Experiment | *P. syringae* strain | Treatments | Figure(s) |
| --- | --- | --- | --- |
| Phage depletion experiment 1 | PT-23 | Microbiome, microbiome + phages, phages | 2,3 |
| LPS challenge | PT-23 | 0.3, 0.6, 1.3, 2.5, 5 EU pure *E. coli* LPS | S1 |
| Phage depletion experiment 2 | DC3000 | Microbiome, microbiome + phages, phages | 2,3 |
| Local adaptation transplant experiment | DC3000 | Microbiome, microbiome + phages, microbiome + phages from neighboring plant, microbiome + phages from different plant species, microbiome + phages from different field | 4, 5 |

**Figure 2.**
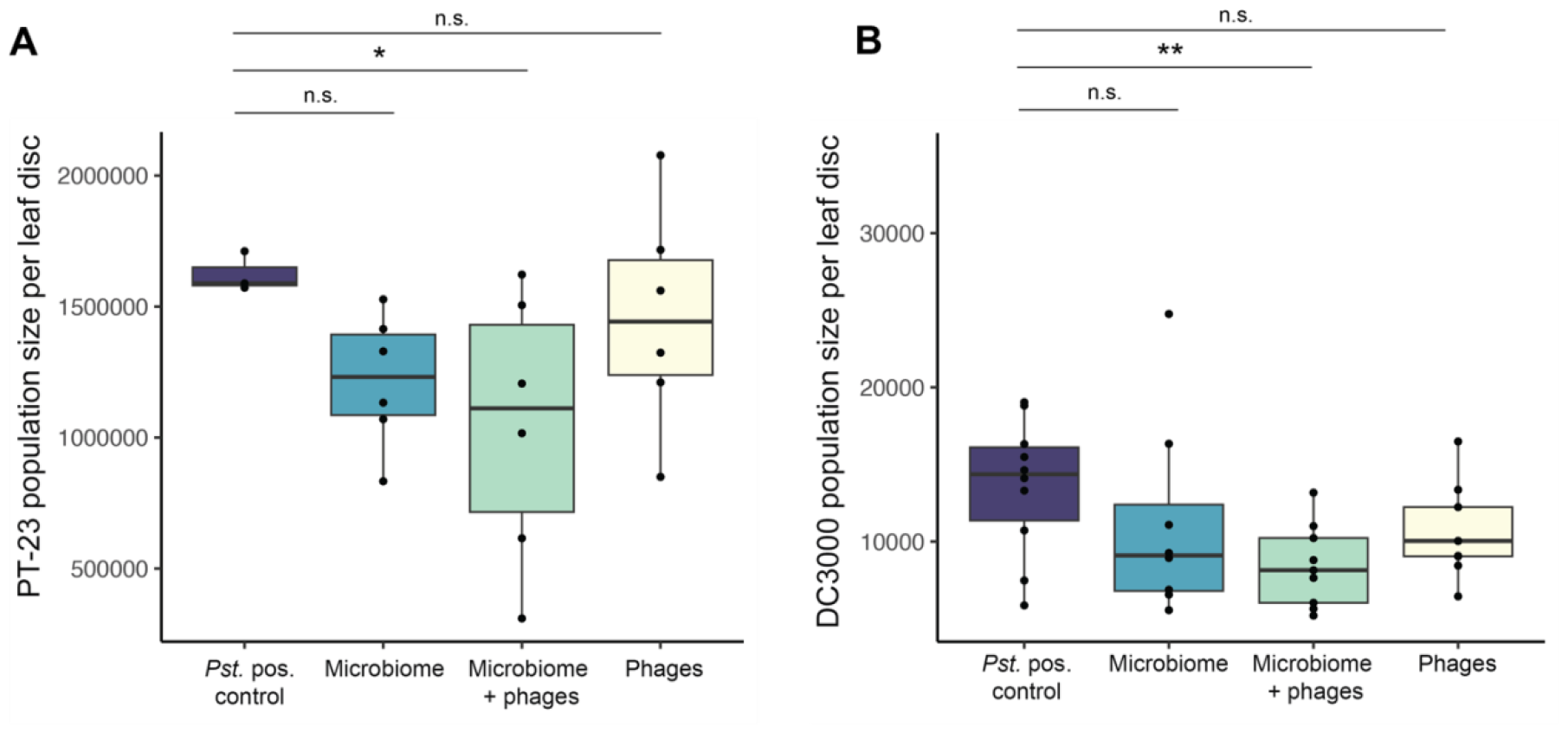
Supplemental microbiome and phages reduce *Pseudomonas syringae* colonization of tomato leaves in two independent harvest seasons. **(a)** Population sizes of *Pseudomonas syringae* pv. tomato strain PT-23 on plants treated with microbial and/or phage communities collected from an agricultural field plot in 2018. **(b)** Population sizes of *Pseudomonas syringae* pv. tomato strain DC3000 on plants treated with microbial and/or phage communities collected from an agricultural field plot in 2021.

To validate and generalize the results of the first experiment, we collected tomato leaves in a subsequent harvest season and repeated the phage depletion process. We sprayed tomato plants with microbial communities as before, but this time we challenged the leaves with a more generalist strain of the same species, *P. syringae* pv. tomato strain DC3000. Compared to the more specialized strain PT-23, DC3000 reached much lower population sizes on tomato plants. Control plants that were treated with only DC3000 harbored 10^2^-10^4^ copies of the 16S gene per 6-mm leaf disc, compared to 10^6^ copies per disc in control plants infected with PT-23. As before, *P. syringae* colonization was significantly reduced in plants treated with the ‘microbiome + phages’ inocula (t = 2.81, df = 35, p = 0.008), while neither component was individually protective (Figure **2b**).

### Direct phage infection of pathogen does not explain the observed protective effect

We first considered the possibility that phage communities aided in microbiome-mediated protection because they contained, by chance, phages capable of directly infecting *P. syringae*. While this seemed unlikely given that phage communities were not significantly protective on their own, it is possible that free phages without hosts decayed rapidly in the week between microbiome treatment and pathogen challenge. With their hosts present in the ‘microbiome + phages’ treatment, phages would be able to maintain sufficient population sizes to subsequently infect *P. syringae* and limit its colonization.

To test this possibility, we co-cultured each phage community with each strain of *P. syringae* to amplify any infective phages if they were present, then plated the co-culture filtrate onto bacterial lawns of *P. syringae*. Only one of the six phage communities in the first harvest season, F18.5, produced plaques on the *P. syringae* plates. This phage community was associated with unusually low *P. syringae* growth in the plant experiment as well, suggesting that a *P. syringae*-targeting phage in this community may have directly reduced *P. syringae* colonization *in planta* (Figure **3a**). Of note, the statistical effect of the ‘microbiome + phages’ treatment did not qualitatively change when the data were reanalyzed to exclude bacteria and phage communities from site F18.5 (t = 2.73, df = 14, p = 0.016), indicating that the protective effect was not solely driven by direct infection by a single phage.

**Figure 3.**
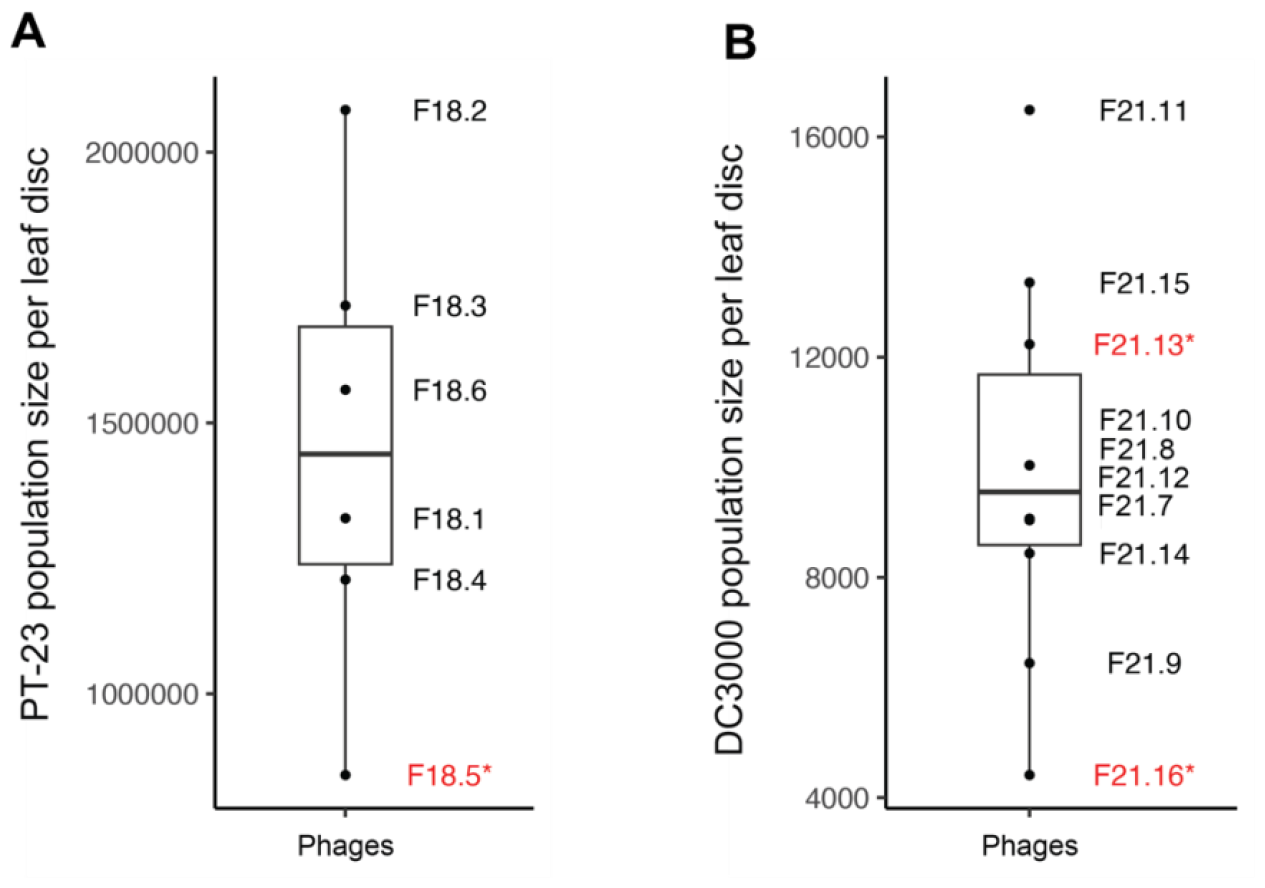
Identifying *Pseudomonas-syringae*-targeting phage activity in field-collected phage communities. **(a)** Population sizes of *Pseudomonas syringae* pv. tomato strain PT-23 on plants treated with phage communities collected from an agricultural field plot in 2018. **(b)** Population sizes of *Pseudomonas syringae* pv. tomato strain DC3000 on plants treated with phage communities collected from an agricultural field plot in 2021. Phage communities in red with an asterisk indicate plaque-forming activity on agar plates of the respective strain of *Pseudomonas syringae*.

In the second harvest season, two out of ten phage communities contained phages capable of infecting *P. syringae*, including the community associated with the lowest *P. syringae* colonization in the plant experiment (Figure **3b**). Again, the effect of the ‘microbiome + phages’ inocula remained significant when these field sources were excluded from analysis (t = 2.31, df = 35, p = 0.029). These results indicate that although direct phage infection of *P. syringae* may have occurred occasionally in our experiments, it was not widespread and could not fully account for the effect of phages on microbiome-mediated protection.

### Lipopolysaccharide accumulation in filtrate does not explain the observed protective effect

We next considered the possibility that the viral isolation process may have accumulated other particles in addition to phages. In particular, ultrafiltration has the side effect of concentrating lipopolysaccharide (LPS), a component of the outer membranes of Gram-negative bacteria (54). If LPS molecules in the phage filtrates trigger a plant immune response, such an effect could be mistakenly attributed to a protective effect of phages. We note that this explanation seemed unlikely, as it expects the phage communities to limit *P. syringae* growth even in the absence of their respective microbial communities, which was not the case in our plant experiments. Nevertheless, we sought to exclude this possibility by quantifying the LPS concentration in the phage filtrates.

LPS concentrations in our phage filtrates ranged from 1-3.5 endotoxin units per milliliter (EU/mL), which fall on the low end of the typical range of reported LPS levels of phage preparations in other studies (54–56). When we sprayed tomato plants with pure bacterial LPS in this range of concentrations, we found no relationship between the concentration of pure LPS applied to the plant leaves and the subsequent colonization of *P. syringae* (t = 0.627, df = 3, p = 0.575, Figure **S1a**). Furthermore, plants that were treated with LPS (at any concentration) prior to the pathogen challenge had similar infection outcomes to plants that were not (t = 0.262, df = 6, p = 0.802). Finally, there was no relationship between the LPS concentrations of the phage filtrates and their ability to limit *P. syringae* colonization (Figure **S1b**, t = -0.48, df = 4, p = 0.656).

### Phage contributions to plant pathogen defense depend on microbiome identity

We next explored whether the ability of phages to limit *P. syringae* colonization depended on interactions with microbial hosts; for example, by altering competitive dynamics or species diversity within the bacterial microbiome. We sprayed microbial and phage communities from tomato plants in the field onto juvenile tomato plants in the lab. A set of six focal microbial communities were paired in turn with (i) phage communities isolated from the same tomato plant in the field, as was the case for all preceding experiments (‘sympatric’), (ii) phages isolated from a different tomato plant in the same field (‘allopatric neighbor’), (iii) phages isolated from an American black nightshade plant in the same field (‘allopatric species’), (iv) phages isolated from a tomato plant grown in a different field (‘allopatric distance’) (Figure **4a**).

**Figure 4.**
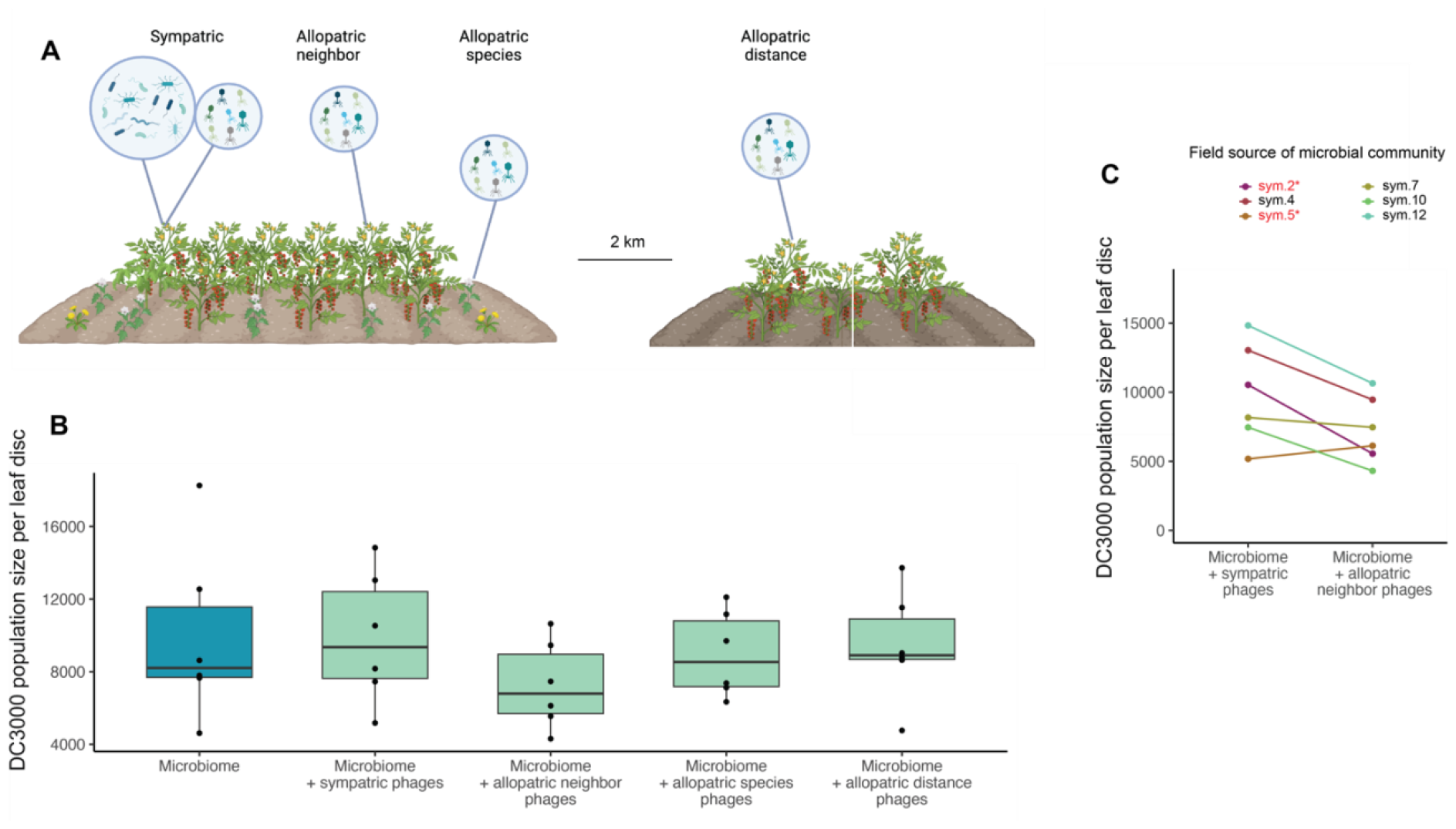
Phages from a neighboring plant more effectively reduce *Pseudomonas syringae* colonization of tomato leaves than phages from the same plant as the supplemental microbiome. **(a)** Diagram showing how phage communities were collected to generate the ‘allopatric neighbor’, ‘allopatric species’, and ‘allopatric distance’ treatments. **(b)** Population sizes of *Pseudomonas syringae* pv. tomato strain DC3000 on plants treated with microbial and/or phage communities collected from an agricultural field plot in 2021. **(c)** Population sizes of *Pseudomonas syringae* pv. tomato strain DC3000 on plants treated with microbial and ‘sympatric’ or ‘allopatric neighbor’ phage communities, colored by the identity of the microbial community. Red text with an asterisk indicates plaque-forming activity on agar plates of the respective strain of *Pseudomonas syringae*.

Bacteria paired with phages from neighboring tomato plants were consistently more protective against *P. syringae* DC3000 than bacteria paired with phages from their own plant (paired t-test, t = 2.82, p = 0.037, Figure **4b**). The sole exception to this pattern, phage source ‘sym5’, contained a *P. syringae*-targeting phage, suggesting that it deviated from the pattern because it substantially reduced *P. syringae* colonization no matter which bacterial community it was paired with (Figure **4c**). The difference between phages from neighboring plants and phages from the same plant remained significant when excluding ‘sym5’ and the other field source with a *P. syringae* phage (paired t-test, t = 3.80, p = 0.032). The two more distant forms of allopatry, phages from a different plant species and phages isolated from a different field, were no different than sympatric phages with respect to protectiveness (p>0.05).

To understand why phages from neighboring plants enhanced microbiome-mediated protection more than phages from the same plant as the bacteria, we sequenced the 16S V4 amplicon of communities from the ‘microbiome only’ transplant, the ‘microbiome + sympatric phages’ transplant, and the ‘microbiome + allopatric neighbor phages’ transplant. There were no differences in bacterial alpha diversity among the three treatments (Figure **5a**). However, compared to the ‘microbiome only’ controls, transplanting with sympatric phages shifted the final composition of each bacterial community more than transplanting with allopatric phages (Figure **5b**, paired t = 3.85, df = 5, p = 0.012). This suggested that phages were on average locally adapted to their bacterial hosts and that more bacterial infections took place in the sympatric pairing. We hypothesized that sympatric phages reduced bacterial population sizes to the point of killing off many protective bacteria, whereas allopatric phages may have occupied a middle ground of shifting the bacterial community without overly disturbing it. Using droplet digital PCR, we measured the total abundance of the bacterial community on the leaves. We confirmed that sympatric phages significantly reduced bacterial abundance compared to the ‘microbiome only’ transplant (Figure **5c**, t = -2.46, df = 7.71, p = 0.041). Allopatric phages also reduced bacterial abundance compared to the ‘microbiome only’ transplant, but the difference in this case was not significant (t = -1.85, df = 6.64, p = 0.108).

**Figure 5.**
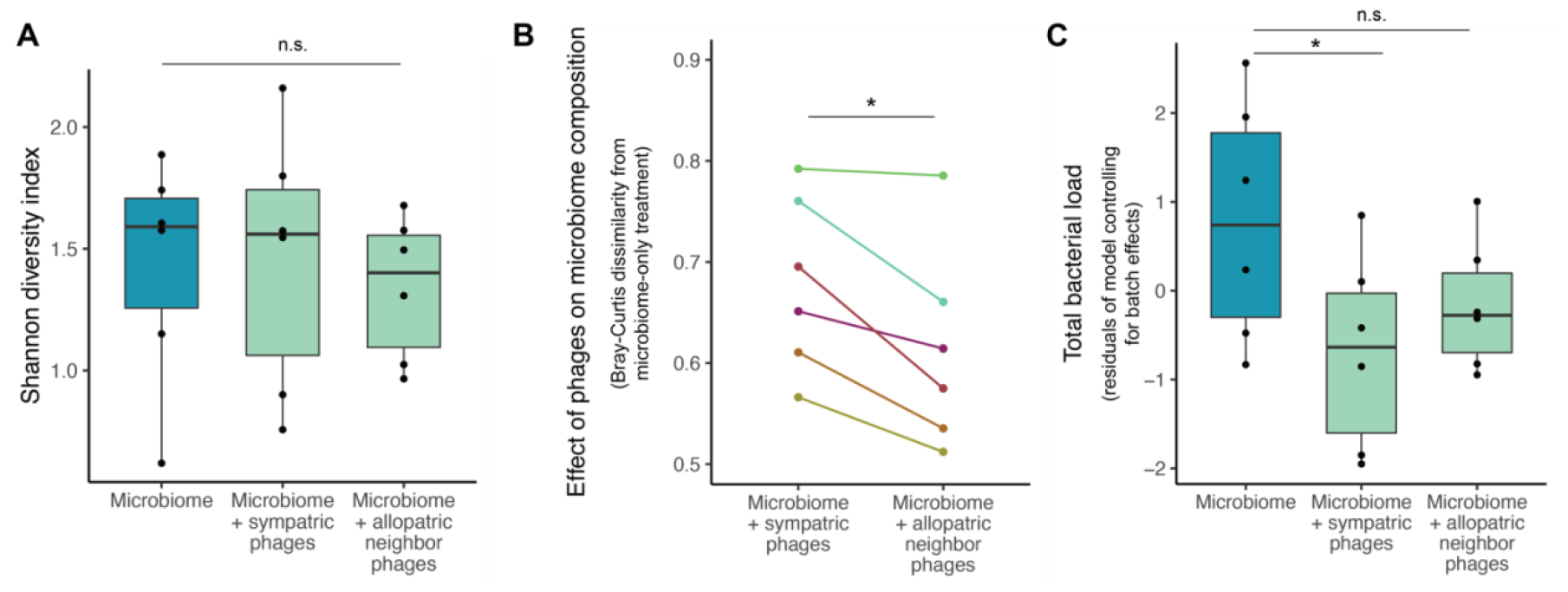
Composition of microbial communities after transplant with or without phages. **(a)** Shannon diversity of microbial communities in each treatment from the sympatric and allopatric transplant experiment. **(b)** Bray-Curtis dissimilarity of microbial communities in each ‘microbiome + phages’ treatment compared to ‘microbiome only’ control plants. Higher values indicate a larger degree of community change. **(c)** Total abundance of microbial communities on leaves after microbiome and phage transplant. For visualization purposes, values are displayed after controlling for row- and column-wise spatial effects in the droplet digital PCR assay.

## DISCUSSION

For many decades, the majority of clinical microbiology research and interventions focused solely on bacterial pathogens. The relatively recent expansion of the microbiome field revealed that pathogens comprise only a small proportion of microorganisms and that commensal microbiome regulation plays an important role in immune health as well (59,60). In a similar vein, though phages have primarily attracted interest in medical microbiology for their ability to infect bacterial pathogens, our work and other recent studies highlight phages of the commensal microbiome as an important component of health and disease (16,61). We found that microbiome and phage communities, when applied to tomato leaves, limited colonization of the bacterial pathogen *Pseudomonas syringae*. This effect persisted across independent field sampling seasons and against two different strains of *P. syringae*.

Phage communities could influence disease susceptibility in several ways (Figure **1**), and we designed our experiments to disentangle these possible mechanisms. We first asked whether the phage communities we had sampled from the field contained, by chance, phages capable of directly infecting and killing *P. syringae*. Several phage communities contained phages that formed plaques on *P. syringae* plates. Some of them were particularly effective at limiting *P. syringae* colonization in the plant experiments as well, while others produced plaques *in vitro* but were not notably protective *in planta*. However, we continued to observe disease-protective effects of the phage treatment when we excluded all phage communities with plaque-forming activity from analysis.

We next explored whether phage communities acted by directly stimulating the plant immune system. Such an effect has been documented in mammals; mice infected with the filamentous phage Pf mount an antiviral response that suppresses antibacterial immunity (23). However, the phage communities in our study only limited *P. syringae* colonization in the presence of the commensal microbiome, suggesting no direct effects on the plants (Figure **2**). Lipopolysaccharide (LPS) accumulation in the phage filtrates was also not sufficient to explain the observed results, as directly applying the corresponding quantities of purified LPS to plant leaves did not recapitulate the protective effect (Figure **S1**).

The observation that phage communities only limited *P. syringae* colonization in the presence of the commensal microbiome suggested that the mechanism of action involved ecological interactions between bacteria and phages. For example, phage lysis of bacterial cells may have altered competitive dynamics between bacteria (6,62) or released bacterial byproducts that were recognized by the plant immune system (24). Temperate phages in the phage filtrates may also have integrated into bacterial genomes and supplied auxiliary metabolic genes or toxins with antimicrobial activity against *P. syringae* (63,64). In an attempt to distinguish between some of these possibilities, we conducted a final experiment that paired microbial communities with either their own phages or phages from increasingly distant sources. Phages consistently had a stronger effect on pathogen protection when they came from a neighboring plant of the same species than when they came directly from the same plant (Figure **4**). Phages from more distant sources, such as a different plant species or a field several kilometers away, were not notably protective, suggesting that the ‘allopatric neighbor’ phages perhaps represented an optimal intermediate.

Amplicon sequencing and droplet digital PCR measurements of the microbial communities suggested that bacterial lysis and community turnover occurred at higher rates when bacteria were paired with their own phages than with neighboring phages. This is consistent with past reports that phages in nature are typically ‘ahead’ of bacteria in coevolution; that is, sympatric phages are typically more infective than allopatric phages (65–67). Given these observations, it appears less likely that phage contributions to pathogen protection were mediated by bacterial products released during lysis, as this would predict the sympatric combinations to be more effective. The possibility that temperate phages carried genes involved in defense seemed less likely as well, as the same set of phage communities with the same gene repertoires was used in both treatments. Like obligately lytic phages, temperate phages have restricted host ranges (68,69), suggesting that they, too, should be more active in sympatric combinations. Instead, a possible explanation for our results is that moderate amounts of phage activity restrict dominant bacterial populations enough to allow other, potentially pathogen-protective bacteria to grow – i.e., frequency-dependent selection or the Kill-the-Winner hypothesis (37–39) – but high levels of phage lysis reduce bacterial populations to the point of actually leaving them more vulnerable to pathogen infection.

The idea that sympatric phage communities overshoot the protective effect by killing protective bacteria naturally raises the evolutionary question of why plant hosts would harbor phage communities that are maladaptive in terms of pathogen defense. One possibility is that phages are less under host control than other taxa and instead respond primarily to selection exerted by their microbial hosts, as suggested from observations that phage communities co-vary closely with microbial communities across hosts (70). Alternatively, endogenous phage communities may be adaptive for occasional or low-level immune challenges (9), but not for the intensity of the pathogen challenge in this study, or they may be particularly impactful during the early stage of microbiome formation that we focused on during our study. Finally, although the microbial and phage inocula in this study were always adjusted to match their relative proportions in the field, the process of recombining them in buffer may have intensified infection dynamics. Bacterial infections of plants, for example, are often facilitated by a wetter environment (71,72), and water droplets have been identified as a potentially important medium for phage diffusion as well (73).

Overall, our data point to an indirect yet important role for phage communities in plant defense against the pathogen *P. syringae*. This phenomenon was not attributable to phage-plant interactions or phage infection of *P. syringae*, as it was largely dependent on the presence and identity of the commensal microbiome. These results suggest an emerging picture of a ‘Goldilocks effect’ in which moderate frequencies of phage lysis are optimal to maintain a protective microbiome. Together, this work highlights a need to consider phages in studies of microbiome-mediated resistance to bacterial pathogens, as they likely have untapped potential for our understanding of dysbiosis and disease.

## MATERIALS AND METHODS

### Microbiome sampling from field

In the first field season (September 2018), leaves were sampled from approximately six-month-old tomato plants at the Student Organic Farm at the University of California, Davis (38°32’20.04” N, 121°44’57.36” W). Leaves were collected from the full range of the height of the plants and placed in a loosely filled bag (35-50 grams of plant material). Bags were transported to the lab and immediately stored at 4°C. Bacteria and phage communities were isolated and frozen within one week of field sampling. These samples were used for the first phage depletion experiment as well as the LPS quantification (Table **1**).

In a subsequent field season (August 2021), leaves were sampled from 16 six-month-old tomato plants at the Student Organic Farm, as well as 6 tomato plants and 6 American black nightshade plants (*Solanum americanum*) at the Vegetable Crops Field at the University of California, Davis (38°32’31.2” N, 121°45’46.8” W). Leaves were transported to the lab on ice, stored at 4°C, and processed within one week of sampling. Of the former group, 10 of these plants were used to generate inocula for the second phage depletion experiment, while the remaining 6 constituted the ‘field allopatric’ treatment in the local adaptation transplant experiment (Table **1**). Of the latter group, the 6 tomato plants were used to generate inocula for the ‘sympatric’ and ‘neighbor allopatric’ treatments, and the 6 American black nightshade plants were used for the ‘species allopatric’ treatment.

### Isolation of microbial and phage communities

A buffer solution of 10 mM MgCl2 was added to each bag to cover the leaves, and the Ziploc bags were submerged in a Brandon M5800 sonicating water bath for 10 minutes to gently dislodge microbial cells from the leaf surface. The resulting leaf wash was passed through a 20 μm membrane to remove plant tissue. The leaf wash filtrate was then vacuum filtered through a 0.2 μm membrane to separate cellular microbes (fungi, bacteria) from viral particles.

To release microbial cells, the 0.2 μm filter was transferred to a sterile tube and sonicated in 10 mM MgCl_2_. Microbial cells were pelleted at 3500 x *g* for 10 minutes, resuspended in 50% glycerol, and stored at -80°C. The filtrate from the 0.2 μm, containing viruses and small molecules, was transferred to an Amicon Ultra-15 filter unit with a 100 kDa molecular weight cutoff. Amicon filters were centrifuged at 4000 x *g* for 25 minutes to isolate and concentrate viruses. The resulting phage fraction was stored at 4°C.

This method of phage isolation and concentration was originally developed to isolate viruses in seawater (74,75) and has been previously adapted to the tomato phyllosphere (40). It separates the majority of lytic phages from their hosts, but lysogenic phages or lytic phages actively infecting a bacterial cell at the moment of filtration are likely to remain in the bacterial fraction, thus resulting in a “phage-depleted” rather than an entirely phage-free microbial community.

### Phage depletion experiments

To measure the effect of phage depletion on microbiome-mediated resistance to pathogen colonization, microbial inocula were applied to plant leaves, either with or without their respective phage communities. Briefly, Early Girl tomato seeds (Eden Brothers) were surface sterilized by immersing seeds in 70% ethanol for 1 minute, then swirling seeds in 10 mL of 6% bleach and 10 mL of 0.2% Tween 20 for 20 minutes. Seeds were placed in a petri dish containing 0.8% water agar, then covered and incubated at 21°C in the dark. After germination, plates were maintained in a growth chamber at 24°C and 70% humidity with a 15h day:9h night light cycle. Nine days after planting, seedlings were transferred to pots containing autoclaved potting medium (Profile Porous Ceramic Greens Grade soil amendment, Sierra Pacific Turf Supply). Pots were spatially randomized with respect to treatment for the duration of the experiment.

Three weeks after planting, leaves were sprayed with microbial communities from the field. Microbiome and phage inocula were each standardized to a fixed quantity of plant material from the field so that phage-host ratios in the inocula were the same as those in their natural environment. Inocula were prepared in 4 mL of 10 mM MgCl_2_, with 0.04 μL of the surfactant Silwet L-77 added to facilitate microbial adhesion to the leaf surface. Leaves were sprayed from all angles using a 15 mL conical tube fitted with a spray cap.

Four weeks after planting (one week after microbiome inoculation), leaves were challenged with *Pseudomonas syringae* pv tomato. Two different strains of *P. syringae* were used in separate experiments: DC3000, a model plant pathogen that infects *Arabidopsis thaliana* as well as tomato plants, and PT23, a closely related pathovar that is more specialized to tomato plants (76). In both cases, an overnight culture of *P. syringae* was pelleted and diluted to an optical density (OD_600_) of 0.0002. The resulting microbial suspension was infiltrated into the abaxial side of the leaf using a blunt-end syringe. At 24 hours post-infection, three hole punches (6-mm diameter) were taken from each leaf. Leaf discs were homogenized in 1 mL 10 mM MgCl_2_ in a FastPrep-24 5G sample disruption instrument at 4.0 m/s for 40 seconds and stored at -20°C for molecular analysis.

Healthy leaves (not challenged with *P. syringae*) were collected, suspended in 10 mMgCl_2,_ and sonicated, pelleted, and frozen as described above to isolate commensal microbiota. We verified that these sequences would not be contaminated by spillover of *P. syringae* from nearby leaves using droplet digital PCR. The assay detected 0-15 copies of the *Pseudomonas* sequence per 6-mm leaf disc (in comparison to ∼10,000 per leaf disc in infected leaves), comparable to baseline levels of *Pseudomonas* probe activity in leaves from plants that were never challenged with *P. syringae* at all (Table **S1**).

### Lipopolysaccharide quantification

The Pierce Chromogenic Endotoxin Quant Kit (ThermoFisher Scientific Cat. #A39553) was used to measure lipopolysaccharide (LPS) concentrations in the phage filtrates. Briefly, amebocyte lysate that binds to LPS was added to samples and incubated at 37°C for 30 minutes. A chromogenic substrate that reacts with the amebocyte proenzyme was added and incubated at 37°C for 6 minutes. Acetic acid was added to stop the reaction, and optical density values were recorded at 405 nm. Absorbance values were adjusted to blanks and LPS concentrations of samples were calculated based on the standard curve.

To assess whether LPS accumulation was responsible for the observed effect of phage on disease outcomes *in planta*, the leaves of three-week-old tomato plants were sprayed with either phage communities or varying concentrations of pure LPS from *Escherichia coli*. Leaves were challenged one week after spraying with *P. syringae* and harvested as described above.

### Reciprocal transplant experiment

Tomato plants were grown and maintained in the growth chamber as described above. The following inocula were applied to three-week-old plants: 1) microbiome only, 2) microbiome with sympatric phages (isolated from the same plant), 3) microbiome with allopatric phages (isolated from a neighboring tomato plant), 4) microbiome with allopatric phages (isolated from a different plant species, American black nightshade, in the same field), and 5) microbiome with allopatric phages (isolated from tomato plants in a different field, approximately 2 km away). One week after microbiome inoculation, leaves were challenged with *P. syringae* and harvested as described above.

### Test for direct phage infectivity

To test whether phage communities contained any phages capable of directly infecting *P. syringae*, co-cultures of *P. syringae* and 100 μL of each phage fraction were incubated in King’s B Broth for 24 hours at 28°C. The resulting overnight culture was passed through a 0.2 μm filter to isolate any phages that might have amplified in the presence of *P. syringae*. Next, 200 μL of *P. syringae* overnight culture was mixed with 2 mL of King’s B Broth supplemented with 0.6% agar. The soft agar mixture was spread evenly onto petri dishes and allowed to dry, then 30 μL of filtrate from the overnight culture was pipetted on top. Plates were incubated at 28°C and monitored daily for signs of bacterial lysis.

### Quantification and analysis of phytopathogen population sizes

*P. syringae* population sizes on leaves were measured using the Bio-Rad QX200™ Droplet Digital PCR system. Reaction mixtures were prepared using 11 μL Supermix for Probes (no dUTP), 1.1 μL probe (forward 5’-ACTTTAAGTTGGGAGGAAGGG-3’; reverse 5’-ACACAGGAAATTCCACCACCC-3’, probe TGCCAGCAGCCGCGG), 5 μL template, and 4.9 μl molecular grade water. Samples were randomized on the plates, with a no-template control in the last well of each column. The following cycling conditions were used for amplification: 95°C for 10 minutes; 40 cycles of the following: (94°C for 30 seconds, 60°C for 1 minute, 72°C for 1 minute); 98°C for 10 minutes; hold at 12°C. Droplet thresholds were set by column based on the fluorescence values in the range of the negative control.

The ability of phage depletion to limit *P. syringae* colonization was assessed using linear regression. For each experiment, the dependent variable was the copies of *P. syringae* detected per standardized disc of infected leaf tissue, which was approximately normally distributed and had similar variance among treatments. The independent variable was the treatment, with the positive control (plants sprayed with a sterile buffer solution in place of a supplemental microbiome prior to *P. syringae* challenge) as the reference level. In the experiment testing the effect of pure LPS on *P. syringae* colonization, the dependent variable was again the number of copies of *P. syringae* per leaf disc and the independent variable was the concentration of LPS (endotoxin units per milliliter). In the reciprocal transplant experiment, the effect of different phage communities on *P. syringae* was assessed using a paired t-test, with pairs of samples linked if they were treated with the same microbial community but different phage communities. This model structure controlled for the fact that microbial communities from different field sources had different intrinsic resistance to *P. syringae* colonization. It tested whether the identity of the phage community (sympatric or allopatric) affected the intrinsic resistance levels of the various microbial communities in the same direction.

### DNA extraction and sequencing

DNA was extracted from leaf wash filtrate from the following treatments: microbial inocula from the field prior to transplant, microbiomes transplanted in the absence of phages, microbiomes with their sympatric phages, and microbiomes transplanted with phages isolated from a neighboring tomato plant in the field. Extractions were performed using the DNeasy PowerSoil kit. Libraries were prepared by amplifying the V4 region of the 16S rRNA gene. Libraries were amplified, cleaned, and sequenced alongside DNA extraction controls and PCR controls on the Illumina MiSeq platform at Microbiome Insights (Vancouver, BC, CAN).

Reads were analyzed using the recommended DADA2 workflow (77) to infer amplicon sequencing variants. Forward reads were truncated at position 240 and reverse reads were truncated at position 140. Reads were filtered to allow 2 or fewer expected errors on the forward reads and 5 or fewer expected errors on the reverse reads. Taxonomy was assigned using the SILVA database (78) and all sequences mapping to plant chloroplast or mitochondria were removed. To compare the effect of phage communities on microbiome composition, a paired t-test was again used to control for the variation among microbial communities from different field sources to phage treatment. The dependent variable was the Bray-Curtis distance between the final microbiome composition at harvest when it was transplanted with phages versus without.

### Quantification of total bacterial abundance

As the amplicon sequencing data revealed high concentrations of chloroplast DNA in our leaf wash samples, we reasoned that accurately quantifying total bacterial abundance would require an additional effort to exclude host DNA. We selected primers to target a region of the 16S rRNA gene that is conserved among bacteria, but has several mismatches with plant chloroplast and mitochondrial 16S rRNA sequences (57). We prepared reaction mixtures for the Bio-Rad QX200™ Droplet Digital PCR system using 11 μL EvaGreen Supermix, 0.22 μL of each primer (forward 5’ - AACMGGATTAGATACCCKG – 3’; reverse 5’ – GACGGGCGGTGWGTRCA – 3’), 5 μL template, and 5.56 μL molecular grade water. Samples were randomized on the plates, with a no-template control in the last well of each column. The following cycling conditions were used for amplification: 95°C for 10 minutes; 40 cycles of the following: (95°C for 30 seconds, 55.4°C for 30 seconds, 60°C for 2 minutes); 4°C for 5 minutes; 90°C for 5 minutes; hold at 4°C. Droplet thresholds were set by column based on the fluorescence values in the range of the negative control. We used known mixtures of bacterial culture and pure extracted chloroplasts to verify that this primer set reduced chloroplast amplification by 90-95% compared to standard 16S V4 primers, while still amplifying the majority of bacteria detected by the standard primers (Figure **S2**).

## ACKNOWLEDGMENTS

This work was supported by an NSF Graduate Fellowship (award #1650114 to RD), a Berkeley Graduate Fellowship (awarded to AC), and the Chan-Zuckerberg initiative (awarded to BK). The authors thank the University of California, Davis for access to the Student Farm plots and members of the Koskella lab for constructive comments.

