## Supplementary Materials for "Phages indirectly maintain plant pathogen defense through regulation of the commensal microbiome"

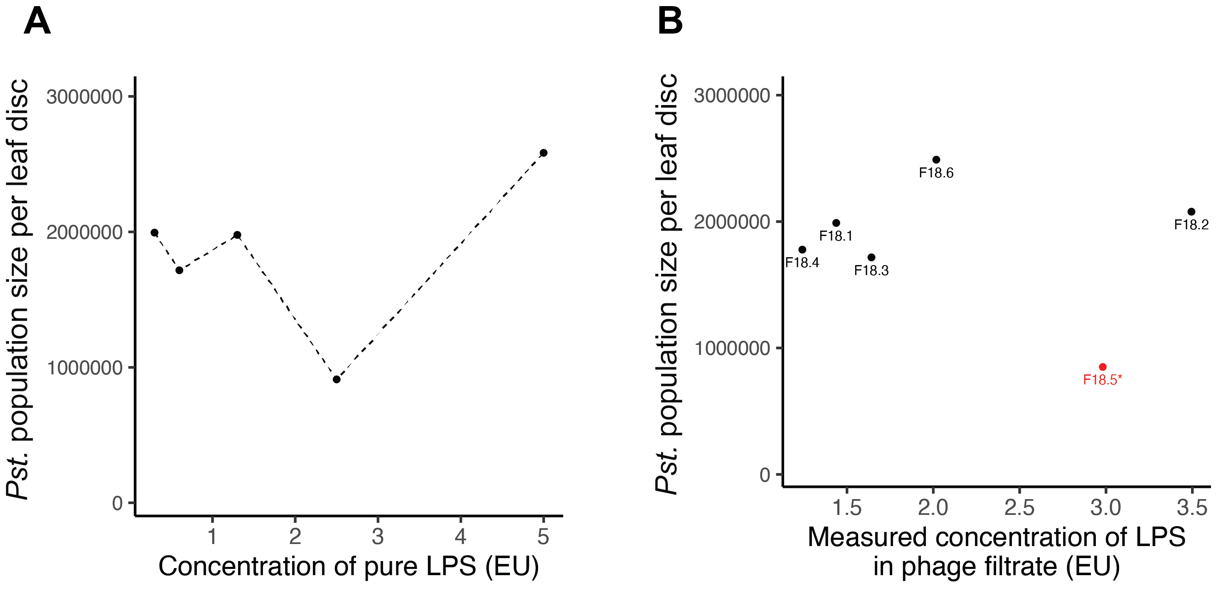

**Figure S1. Relationship between lipopolysaccharide content of treatment and *Pseudomonas syringae* colonization. (a)** Population sizes of *Pseudomonas syringae* pv. tomato strain PT-23 on plants treated with pure bacterial lipopolysaccharide (LPS). **(b)** LPS concentrations of phage filtrates isolated from an agricultural field plot in 2018, and resulting population sizes of *Pseudomonas syringae* pv. tomato strain PT-23 on plants treated with those phage communities. Red text with an asterisk indicates plaque-forming activity on agar plates of the respective strain of *Pseudomonas syringae.*

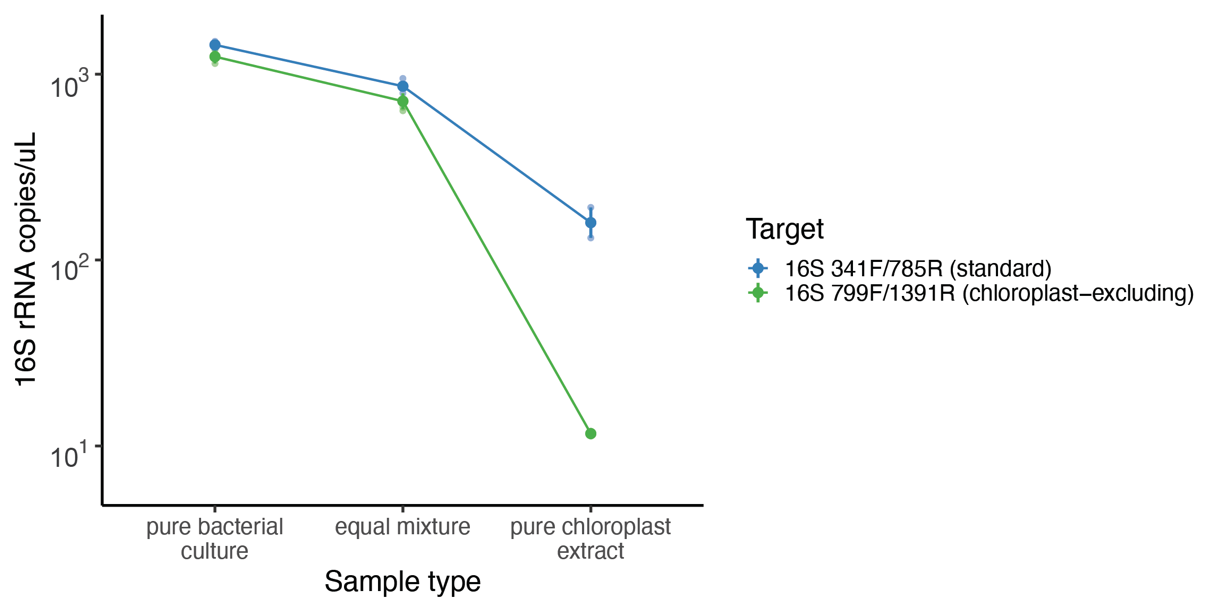

**Figure S2. Validation of chloroplast-excluding primers for bacterial abundance assay.** Droplet digital PCR was conducted on either an overnight bacterial culture of *Pseudomonas putida*, chloroplasts isolated from homogenized tomato leaves by differential centrifugation, or an equal mixture by volume of bacteria and chloroplasts.

**Table S1. *Pseudomonas* abundance in healthy and infected leaves.**

| **Treatment** | ***Pseudomonas* probe signal in infected leaves** | ***Pseudomonas* probe signal in healthy leaves** |
| --- | --- | --- |
| Plants challenged with *P. syringae* | 29685.71 ± 12741.29 | 17.14 ± 4.59 |
| Plants not challenged with *P. syringae* (negative control) | N/A | 0.75 ± 0.75 |
